# LAND USE AND COVER CHANGE IN HALMAHERA, INDONESIA: What are the main vectors of deforestation that affect the Hongana Manyawa?

**DOI:** 10.64898/2026.08.11.744225

**Authors:** Daniel Fernandes de Figueiredo Silva, Luís Felipe dos Santos Melo, Daniel Cangussu

## Abstract

The global market concentrates extractive pressure on lands held by Indigenous peoples, including peoples living in isolation, for whom free, prior and informed consent cannot be obtained and protection must therefore rest on territorial instruments. Halmahera, Indonesia, holds some of the world’s largest lateritic nickel reserves beneath a lowland rainforest inhabited by the Hongana Manyawa, yet the trajectory of land use and cover change across the island has not been quantified. We characterised land use and cover change over the 17,437 km^2^ island between 2014 and 2024 using MapBiomas time series, and projected a business-as-usual scenario to 2054 with a stochastic cellular-automata model implemented in Dinamica EGO, calibrated with weights of evidence on eight variables describing mining and logging concessions, transport infrastructure, settlements and previous clearing. Forest covered 83.0% of the island in 2014 and 82.1% in 2024; under unchanged policy it falls to 73.7% by 2054, a net loss of 162 thousand ha, or 11.2% of the 2014 baseline, at gross rates of 47,000–51,000 ha per decade. Deforestation probability is highest within 500 m of previous clearing and declines with distance from settlements, cities and mining sites, while proximity to national parks carries a negative weight of evidence. The frontier is self-propagating and spatially predictable, and legally designated territory retains forest within it. Protecting the Hongana Manyawa consequently depends on excluding extractive licensing from the interior forest ahead of the frontier rather than behind it.

## 1. Introduction

The decarbonization of the global energy system has displaced extractive pressure onto territories historically marginal to industrialization: more than half of the world’s energy-transition mineral projects are located on or adjacent to lands held by Indigenous or peasant peoples (Owen et al., 2023). Halmahera, the largest island of North Maluku, Indonesia, is an example of this contradiction. This island holds some of the world’s largest lateritic nickel reserves. The deposits are shallow and therefore extractable only through the complete removal of the native forests, while the same substrate sustains a lowland rainforest of high endemism and biogeographic singularity (Monk; de Fretes; Reksodiharjo-Lilley, 1997). Since the 2020 revision of the Mineral and Coal Mining Law and the consolidation of the national downstreaming agenda, the island has repositioned itself as an industrial frontier centered around the Weda Bay–IWIP complex, with tree-cover loss in the central and eastern regencies exceeding 80,000 ha between 2001 and 2022 (Hansen et al., 2013; Global Forest Watch data).

Within this frontier live the Hongana Manyawa (“people of the forest”), Tobelo-speaking uncontacted and recently contacted indigenous peoples. They are estimated at several hundred individuals, and they remain in isolation, sustaining a mobile foraging economy across the interior watersheds (Survival International, 2024). Independent assessments report that at least 5,331 ha of forest were cleared inside nickel concessions island-wide, releasing approximately 2.04 Mt CO_2_e (Climate Rights International, 2024), and that concessions overlap a substantial share of the forest used by groups in isolation (Survival International, 2024). The central governance problem follows directly: for peoples in isolation, free, prior and informed consent is unobtainable, so consent-based safeguards are essentially impossible to obtain, and protection must rest on territorial instruments. Yet a spatial analysis to deepen our knowledge of land use and cover change, so we might be able to forsee the future, and what it holds for isolated peoples, such as the Hongana Manyawa, not been produced yet.

To fill this his knowledge gap we characterize land use and cover change (LUCC) across Halmahera over the last decade (2014-2024) from a time series available on MapBiomas platform, furthermore we quantified the advance of the mining and logging concessions, road and settlement frontier towards the pristine native forests within the island and derive indicators of territorial pressure over the protected areas in Halmahera. We projected a business-as-usual (BAU) scenario, of land use and cover change for the next three decades, if policy remains unchanged.

## 2. Methods

We selected eight variables that presently affect the Hongana Manyawa peoples by directly or indirectly increasing the pressure for land use and cover change. All variables were acquired through web-scraping techniques from publicly available Indonesian datasets and pre-processed Python 3.11.

Land-use and cover change was simulated for future scenarios in Dinamica EGO 8 following the stochastic cellular-automata architecture described by Soares-Filho, Cerqueira and Pennachin (2002) and Soares-Filho et al. (2004), implemented as the ten-step calibration–validation–projection workflow of the platform’s LUCC guidebook (Leite-Filho et al., 2020). We simulated the LUCC trajectory in three one-decade time-steps, from 2024 until 2054.

## 3. Results

Our model was able to estimate land-use cover in ten-year steps across the Halmahera Island. For each analyzed year, the total forest and non-forested regions were quantified. The data, as shown in Figure 2, suggests that forests occupied 83.0% of the mapped area in 2014 and 82.1% in 2024, declining across the simulated steps to 79.2% in 2034, 76.4% in 2044 and 73.7% in 2054. The non-forest class followed the inverse trajectory, rising from 16.8% in 2014 to 17.7% in 2024 and reaching 20.6%, 23.4% and 26.1% in the three subsequent decades.

**Figure 1.**
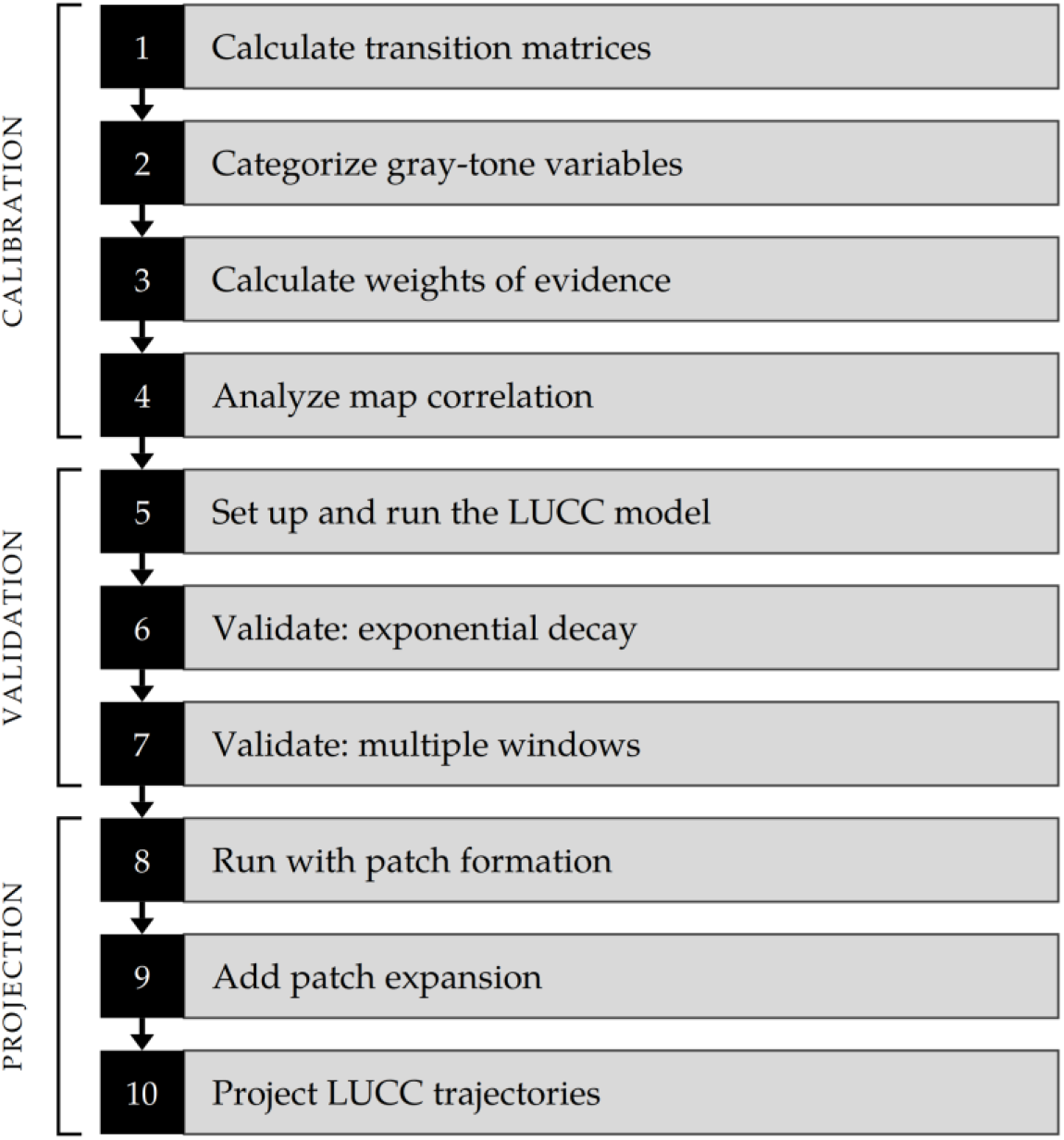
Flowchart of the LUCC model methods. (Leite-Filho et al., 2020)

**Figure 2.**
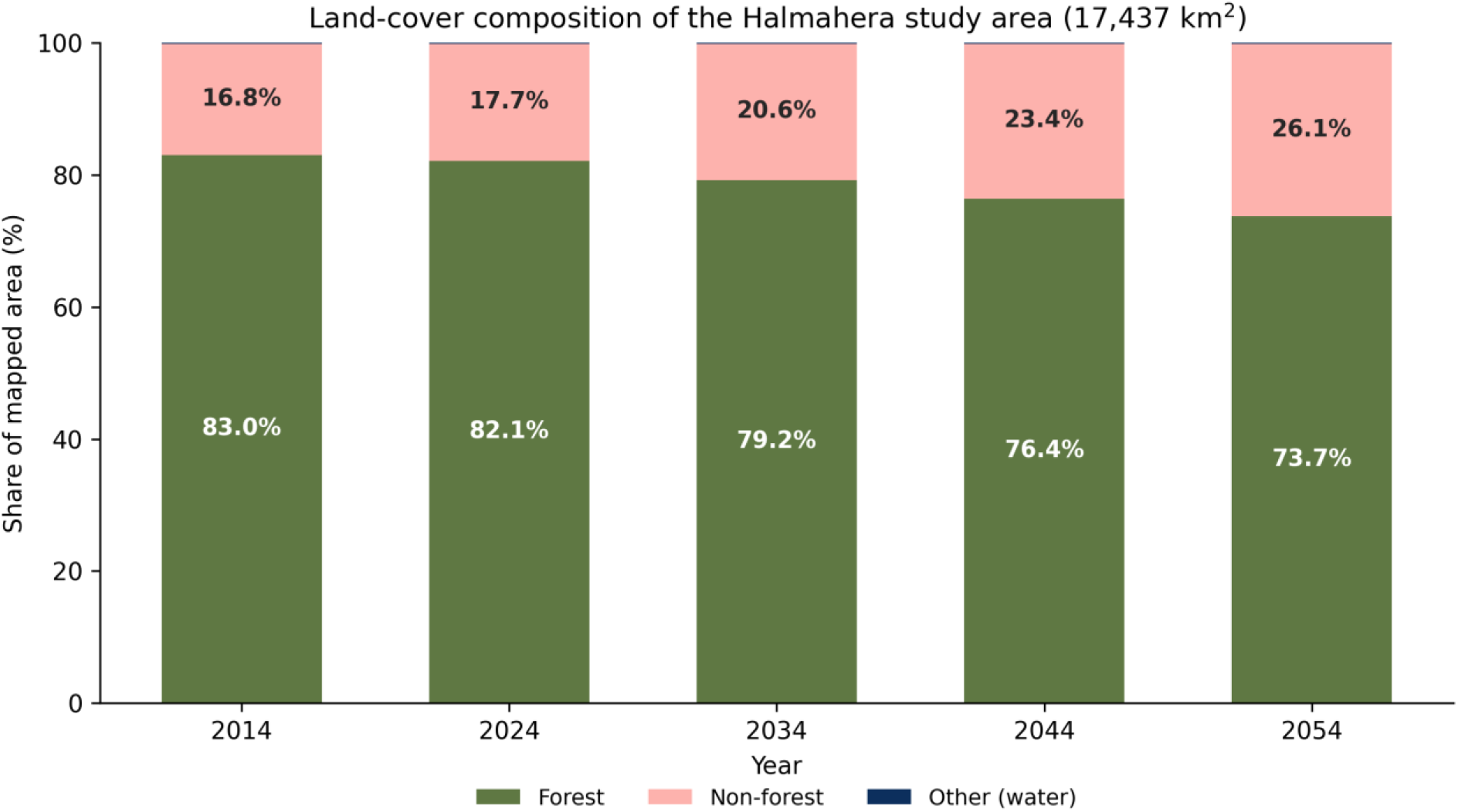
Land use and cover composition of Halmahera Island, Indonesia, from 2014 until 2054.

The weights of evidence (W+) are used to simulate where deforestation is most likely to occur, based on each variable, given the last transitions that took place. Positive values indicate that the variable facilitates the transition. Negative values of W+ suggest that the variable has a repelling effect on the transition. Distance to airports, distance to cities, distance to mining sites, distance to populated places and distance to previous deforestation all show maximum positive W^+^ in the first distance interval, followed by monotonic decline. Distance to National Parks has the opposite effect, indicating maximum repelling effect on deforestation on the first interval, with exponential decrease in W+ (Figure 3).

**Figure 3.**
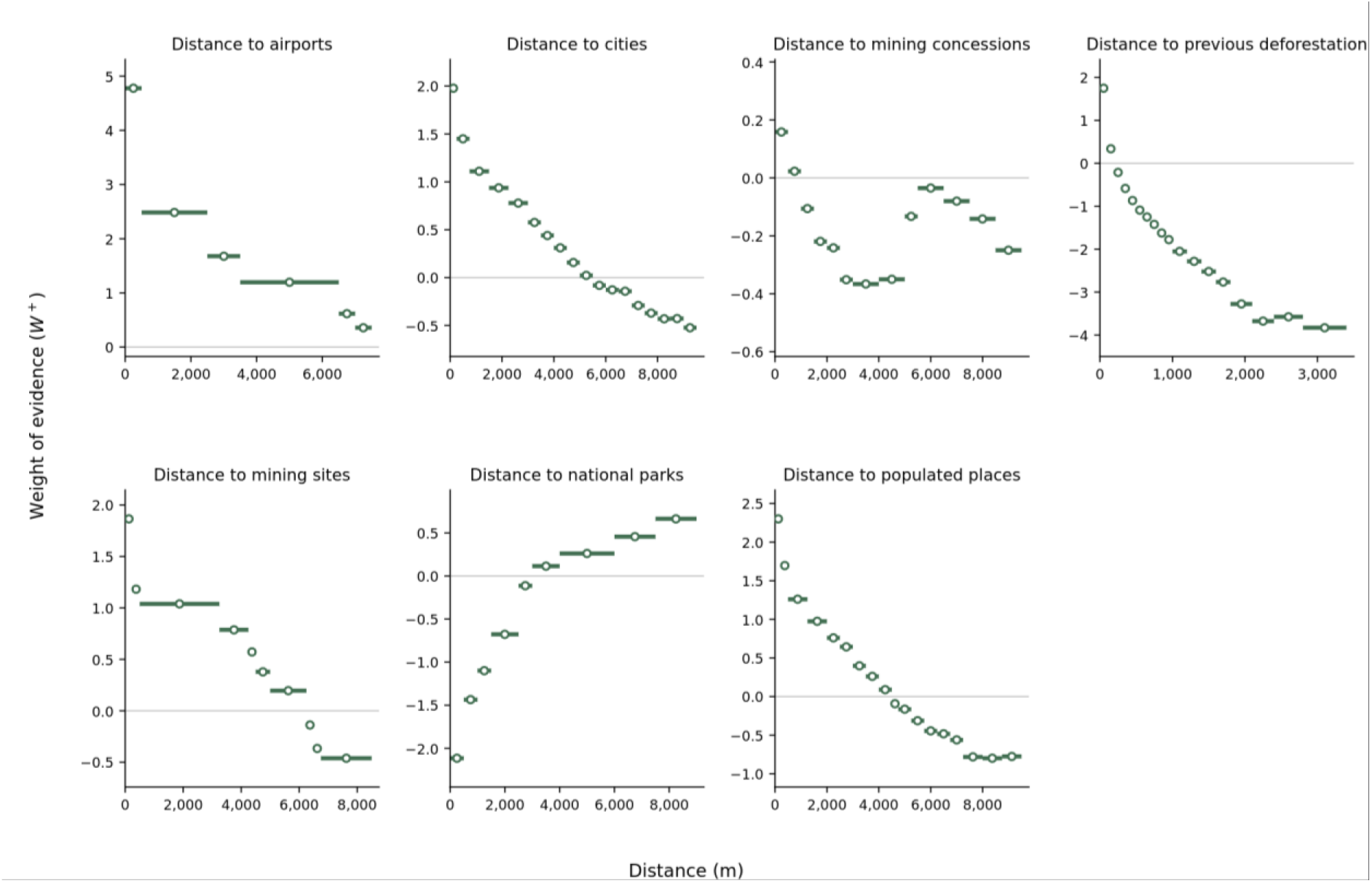
Weights of evidence (W+) for deforestation transitions used in the model. Higher values favor deforestation transitions.

The mean deforestation probability surface for 2024–2054 is presented in Figure 4. Higher probability values concentrate along the coastal margins and in the central and eastern portions of the island, coinciding with the concession polygons. The interior of the national park units and the western and southern uplands retain values near the lower end of the scale.

**Figure 4.**
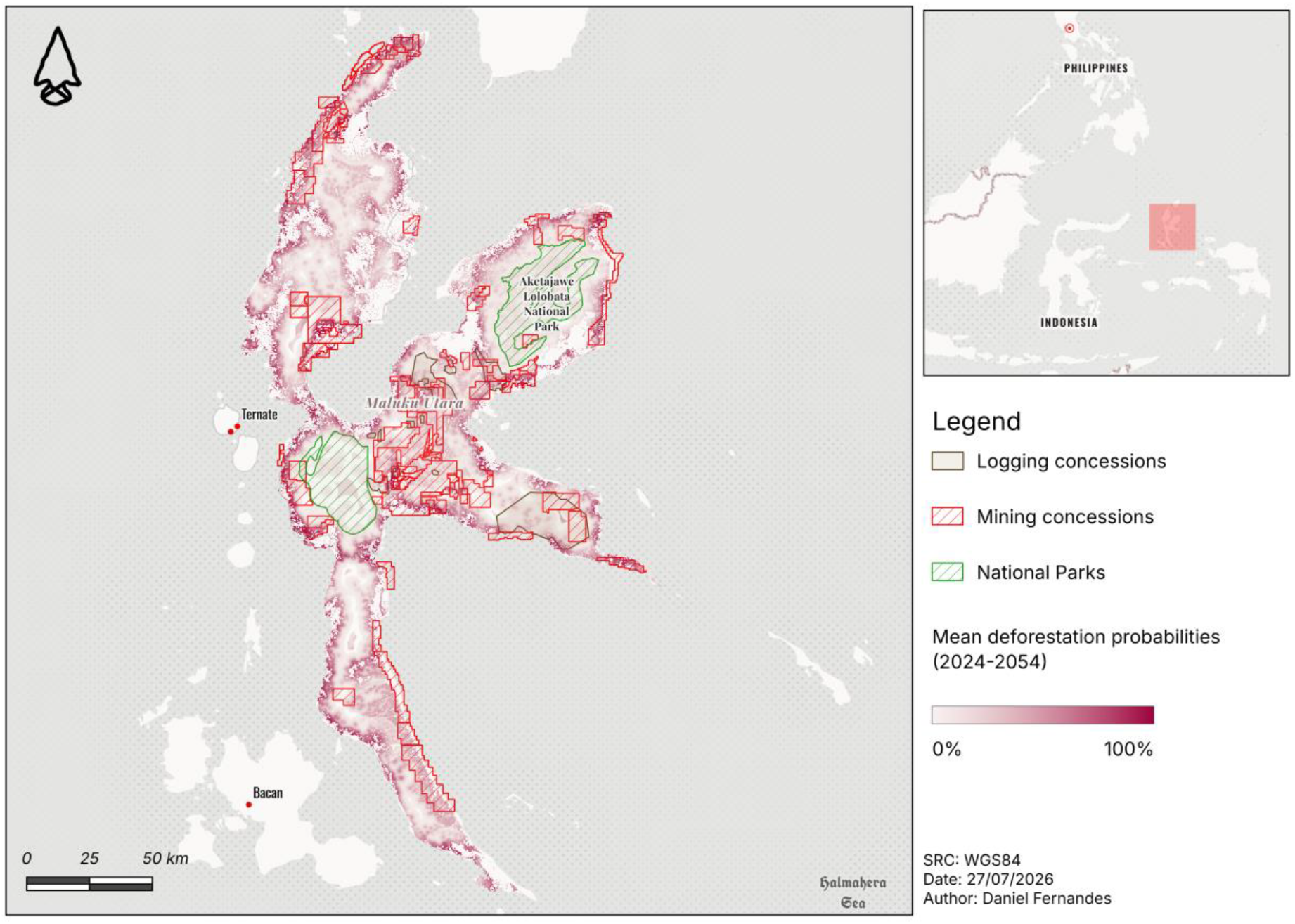
Mean deforestation probabilities between 2024 and 2054.

The contrast between the observed 2024 land-use and cover map with the projected 2054 map is presented in Figure 5. The non-forest class in 2054 appears expanded relative to 2024, principally in the northern and north-eastern portions of the island and along the boundaries of the blocks already classified as non-forest in 2024.

**Figure 5.**
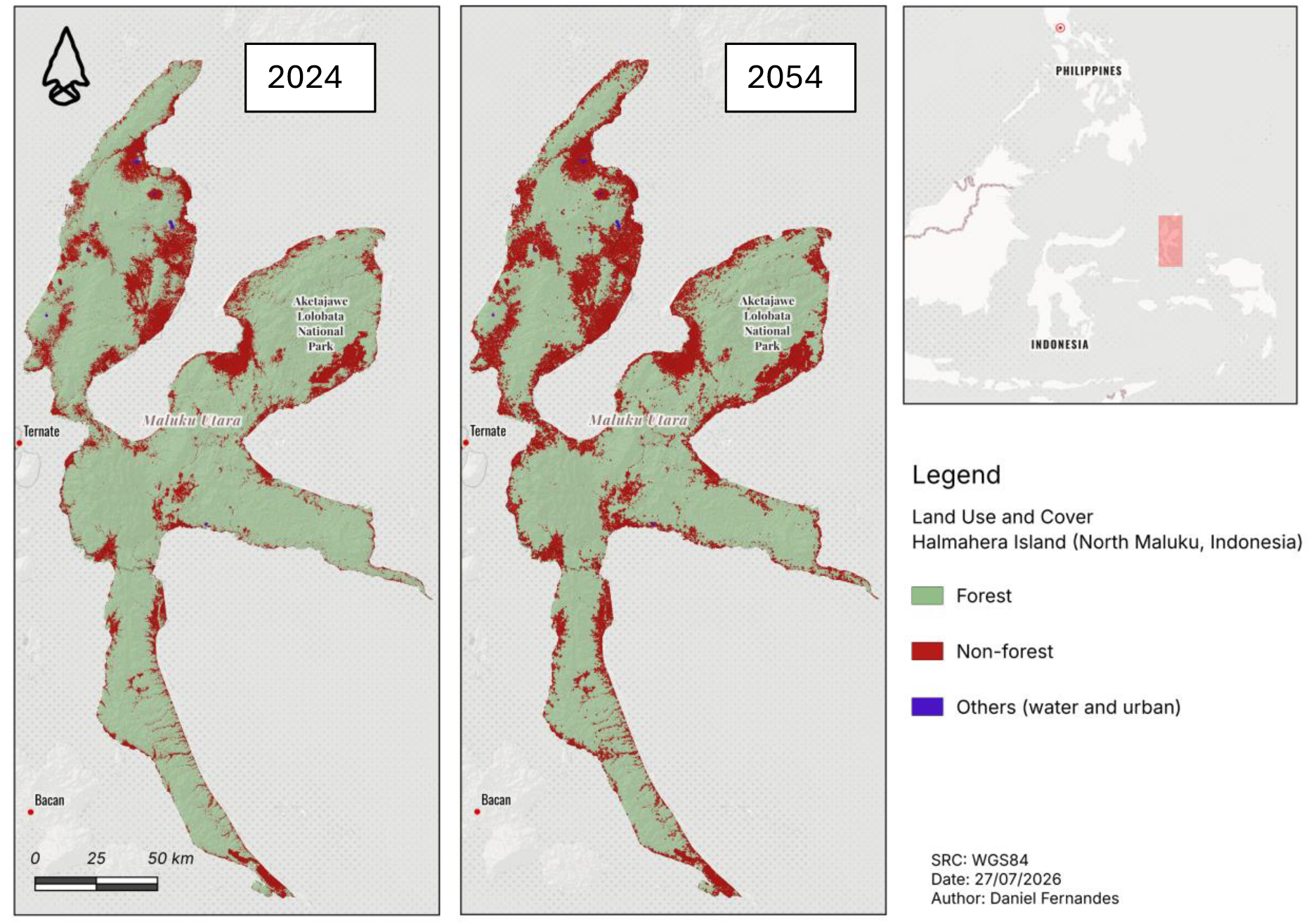
Observed Land Use and Cover (LUC) for 2024 and Projected LUC for 2054.

## 4. Conclusions and Recommendations

Halmahera’s frontier is spatially predictable. Unchanged policy, the Business-As-Usual scenario we projected, indicates the loss of roughly 10% of its forests until 2054. This loss is not diffuse. Deforestion is organized by proximity to previously deforested areas, established settlements and extractive infrastructure. This frontier is dinamic in space, since closeness to deforestation vectors matter, but also in time, since each increment of deforestation favors the next one.

The Hongana Manyawa are surrounded by the de-characterization of their traditional territory. The pressure that reaches peoples in isolation is a function of where infrastructure and concession boundaries are allocated, where Indonesian policy allows. That is why previous, free and informed consent, unobtainable from people who cannot be consulted, is the wrong instrument here. National Parks exhibit strong repelling effect on deforestation. Designating protected areas is therefore the protection instrument available in Halmahera.

Given our results, we suggest the following recommendations:

- Exclude the interior forest used by the Hongana Manyawa from extractive licensing, beginning with the sectors of highest modelled deforestation probability;
- Expand existing protected-area network into the sectors where that forest coincides with the highest modelled deforestation probability.

